# ATM and ATR Influence Meiotic Crossover Formation Through Antagonistic and Overlapping Functions in *C. elegans*

**DOI:** 10.1101/604827

**Authors:** Wei Li, Judith Yanowitz

## Abstract

During meiosis, formation of double-strand breaks (DSBs) and repair by homologous recombination between homologs creates crossovers (COs) that facilitate chromosome segregation. CO formation is tightly regulated to ensure the integrity of this process. The DNA damage response kinases, Ataxia-telangiectasia mutated (ATM) and RAD3-related (ATR) have emerged as key regulators of CO formation in yeast, flies, and mice, influencing DSB formation, repair pathway choice, and cell cycle progression. The molecular networks that ATM and ATR influence during meiosis are still being resolved in other organisms. Here we show that *Caenorhabditis elegans* ATM and ATR homologs, ATM-1 and ATL-1 respectively, act at multiple steps in CO formation to ultimately ensure that COs are formed on all chromosomes. We show a role for ATM-1 in regulating the choice of repair template, biasing use of the homologous chromosome instead of the sister chromatid. Our data suggests a model in which ATM-1 and ATL-1 have antagonistic roles in very early repair processing, but are redundantly required for accumulation of the RAD-51 recombinase at DSB sites. We propose that these features of ATM-1 and ATL-1 ensure both CO formation on all chromosomes and accurate repair of additional DSBs.

**Article Summary:** Crossovers formed during meiosis connect homologs and properly align them for cell division. The central importance of crossovers is underscored by the existence of extensive regulatory processes that ensures the proper execution of these events. This paper explores the evolutionary conserved roles of the central DNA damage response kinases, ATM and ATR, in crossover formation. The authors show that these kinases function together as rheostats to promote timely formation of crossovers on all chromosomes but to limit extensive DNA damage. This work provides a platform for identifying conserved meiotic targets of ATM and ATR that affect fertility across species.

## INTRODUCTION

Crossover recombination, the exchange of DNA between homologous chromosomes, occurs in meiosis I and is a key step to ensure that chromosomes are segregated properly. Many factors contribute to the recombination outcome, including the number of double-strand breaks (DSBs) made during meiosis, their distribution, and the repair pathway chosen following the formation of DSBs. The formation and repair of DSBs and the conversion of some of these into crossovers (COs) is a highly regulated process that must be tightly controlled to ensure the proper subsequent segregation of chromosomes. Mechanisms have evolved to tune DSB numbers to species-specific levels (Gray *et al.* 2013) and also to channel a limited number of DSBs into COs (Martini *et al.* 2006).

The DNA damage response (DDR) kinases ATM/Tel1 (ataxia-telangiectasia mutated) and ATR/Mec1 (ataxia-telangiectasia and RAD-3 related) have emerged as key conserved players in CO homeostasis (reviewed in (MacQueen and Hochwagen 2011; Cooper *et al.* 2014)). ATM mutations lead to severe meiotic defects and infertility in mice, yeast and flies (Barlow *et al.* 1996; Xu *et al.* 1996; Barlow *et al.* 1998). ATM both downregulates Spo11-mediated DSBs (Joyce *et al.* 2011; Lange *et al.* 2011; Zhang *et al.* 2011; Carballo *et al.* 2013; Garcia *et al.* 2015) and affects their distribution (Zhang *et al.* 2011; Anderson *et al.* 2015; Garcia *et al.* 2015). It also biases use of the homolog as a repair template (inter-homolog homologous recombination (IH-HR)) versus the sister chromatid (intersister (IS-HR)). ATR, by contrast, positively regulates DSB formation (Gray *et al.* 2013). Together ATM and ATR establish a regulatory feedback through the phosphorylation of components of the DSB machinery and chromatin axes (Cooper *et al.* 2014). Although yeast ATM and ATR (Tel1 and Mec1, respectively) seem to function antagonistically to enforce DSB homeostasis, the interplay between the two genes and their functions in CO formation are more complex. For example, research in yeast supports that there is a DSB threshold above which Tel1 plays a role (with Pch2 and Mec1) in homolog bias, while with low-abundance DSBs, Tel1 promotes resection (Joshi *et al.* 2015; Mimitou *et al.* 2017). Thus, further insights into the functional consequences of ATM/ATR loss on CO formation through the analysis of additional model systems may help provide new insights into ATM/ATR function.

In *Caenorhabditis elegans*, the orthologs of ATM and ATR, ATM-1 and ATL-1 (ATM-like), are required for genome stability (Jones *et al.* 2012). In addition, more RAD-51 foci were observed in meiotic cells in *atm-1* mutants compared to wild type, a result that was consistent with a conserved role in DSB inhibition (Checchi *et al.* 2014). However, further information about the role of ATM-1 in DSB and CO formation remains unknown. Loss of *atm-1* function has been reported to lead to a mild increase in meiotic nondisjunction suggesting a more complex relationship between DSB formation and CO induction. Because most chromosomes receive a single CO each meiosis (high CO interference) and because only a single CO pathway has been identified in *C. elegans*, we reasoned that the worm offered an opportunity to investigate both conserved features of ATM and ATR functions in meiosis and to dissect out their contributions to unique aspects of CO control that are harder to examine in other systems. Here we show that *C. elegans atm-1* and *atl-1* act at multiple steps in CO formation to ultimately ensure that COs are formed on all chromosomes.

## MATERIALS AND METHODS

### Genetics and worm handling

All strains were grown and maintained at 20° on standard media (Brenner 1974). Mutant strains used in this study were: LGI *atm-1(gk186), rad-54(ok615);* LGII *dsb-2(me96), smc-5(ok2421),); meIs8[unc-119(+) pie-1promoter::gfp::cosa-1];* LGIII *brc-1(tm1145), dpy-18(e364), unc-64(e246), dpy-1(e1), lon-1(e185);* LGIV *spo-11(me44), dsb-1(we11);* LGV *him-5(ok1896), syp-1(me17), atl-1(tm853). atm-1(gk186)* is a deletion allele that removed upstream promoter sequences and half of the 5’ coding sequence; it is a presumptive null. *atl-1(tm853)* is an ~700bp deletion in the coding sequence that is a strong loss-of-function or null. Some strains were provided by the *Caenorhabditis* Genetics Center that is funded by National Institutes of Health - Office of Research Infrastructure Programs (P40 OD010440). Double, triple and quadruple mutants were generated using standard genetic techniques with PCR verification of genotypes and are listed in Supplementary Table S1.

### Immunofluorescence

Adults worms were dissected in 1 X sperm salts with 1 mM levamisole and fixed in 2% paraformaldehyde/1 X PBS for 5 min in a humid chamber. Slides were then freeze-cracked and immersed in 100% ethanol for 2 min followed by 5 seconds in acetone. Slides were then washed in PBSTB (1 X PBS with 0.1% Tween and 0.1% bovine serum albumin (BSA)) and incubated overnight at 4° in primary antibody (rabbit anti-RAD-51, 1:30000, gift from S. Smolikove, rabbit anti-DSB-2, 1:20000, gift from A. Villeneuve) diluted in PBSTB. The next day, slides were washed 3 x PBSTB and incubated in secondary antibody (a-rabbit Alexa 568, 1:2000) for 4 hours at room temperature in the dark. Then slides were washed 2 × 10 minutes, and stained 1 × 10 min with DAPI (10 mg/ml stock diluted 1:50000 in 1 X PBS). Slides were mounted in Prolong Gold with DAPI and put in the dark to dry overnight before imaging.

### Analysis of RAD-51 foci

3D images of the whole germ lines were taken using a Nikon A1r confocal microscope and analyzed using Volocity 3D software (PerkinElmer). For wild type and *atm-1, him-5, dsb-2, atm-1;him-5* and *atm-1;dsb-2* mutants, as well as irradiated and non-irradiated *spo-11* mutants and *atm-1;spo-11*, we divided the pachytene region into six zones and counted RAD-51 foci in every nucleus for a minimum of three germ lines/ genotype. For *rad-54;him-5* mutants and *atm-1;rad-54;him-5* mutants, we counted the RAD-51 foci in late pachytene nuclei for at least three germ lines/ genotype. For analysis of *atl-1* mutants (Figure 7, 8), the transition zone (TZ) was included and the pachytene zones were binned into 3 regions (1+2)/zone (3+4)/zone (5+6) shown in Figure 2A). Three gonad arms were analyzed/ genotype.

### Irradiation

Day 1 adult worms were exposed to γ-irradiation from a ^137^Cs source (Gammacell1000 Elite, Nordion International Inc.). Dosages are described in text. For analysis of diakinesis stage nuclei post-IR, animals were fixed and stained 27 hrs post-IR(Mcclendon *et al.* 2016). For the worms used for RAD-51 staining for time course analysis, we stained them 1, 2, 4, and 8 hrs post-IR.

### Recombination analysis

Recombination rates between *dpy-18* and *unc-64* or *dpy-1* and *lon-1* performed by crossing the marker mutations into the respective genetic background and assaying for. Dpy non-Unc and Unc-nonDpy progeny from *dpy-18 unc-64/+* parents or Lon non-Dpy (Dpy is epistatic to Lon, so cannot be assessed for recombination). More than 1000 progeny for each phenotype were analyzed and the recombination rate was calculated based on prior results(Brenner 1974).

### Analysis of GFP::COSA-1 foci

Day 1 adult worms were dissected in 2 x sperm salts as described above. Slides were immediately freeze-cracked and immersed in 100% ethanol for 10 s and fixed in 2% paraformaldehyde/1 X PBS again for 10 min. Slides were washed 2 × 5 min in PBSTB, stained with DAPI in 1 X PBS for 10 min followed by one wash with PBSTB for 5 min. Slides were mounted in Prolong Gold with DAPI.

Images were acquired and analyzed as described above. GFP::COSA-1 foci in late pachytene nuclei were counted in at least 5 germ lines/ genotype in the late pachytene.

### Data Availability

Strains and plasmids are available upon request. The authors affirm that all data necessary for confirming the conclusions of the article are present within the article, figures, and tables.

## RESULTS

### ATM-1 helps to promote crossover formation

Meiotic DSBs are catalyzed by the conserved topoisomerase SPO11. SPO11 activity is regulated by accessory factors that influence the timing, placement, and extent of DSB formation. ATM influences DSB formation in species as diverged as yeast and mice. In worms, *atm-1* mutant animals are homozygous viable and morphologically wild-type, but variably have offspring with reduced viability and fecundity (Jones *et al.* 2012). In the germ line, an increased number of RAD-51 foci in *atm-1* mutant worms has been interpreted to support a widely conserved role in DSB formation (Checchi *et al.* 2014). In *C. elegans*, mutations in several of the SPO-11 accessory factors, including *him-5* and *dsb-2*, lead to a partial impairment in DSB formation, a subsequent decrease in RAD-51 foci, and a lack of COs on a subset of chromosomes (Meneely *et al.* 2012; Rosu *et al.* 2013). Since *atm-1* mutants have been shown to exhibit an excess of RAD-51 foci that could reflect an excess of DSBs, we wanted to address how SPO-11 accessory factors and ATM-1 interact to impact CO formation. To test this, we constructed *atm-1;him-5* and *atm-1;dsb-2* double mutants and examined bivalent formation in single and double mutants by whole mount fixation and DAPI-staining.

The *atm-1;him-5* and *atm-1;dsb-2* double mutants contained significantly fewer bivalent chromosomes compared to *him-5* and *dsb-2* single mutants (Figure 1A, 1B, P<0.01). The 6 DAPI bodies observed in almost all wild-type germ cells correspond to the 6 bivalents formed between each pair of homologous chromosomes. In *him-5* and *dsb-2* single mutants, the number of DAPI bodies is increased since the non-exchange chromosomes separate from one another into discrete masses, or univalents (Figure 1A, 1B and (Meneely *et al.* 2012; Rosu *et al.* 2013)). The *atm-1* single mutant showed ~5% of nuclei with fewer than 6 DAPI bodies (Figure 1A, 1B). A subset of these may reflect whole chromosome fusions as the result of DNA repair defects (Jones *et al.* 2012). The formation of X:autosome fusions could explain the appearance of heritable, high frequency HIM (high incidence of males) lines in *atm-1* mutants (Jones *et al.* 2012). Surprisingly, close to 10% of *atm-1* mutant nuclei had 7 DAPI bodies.

**Figure 1.**
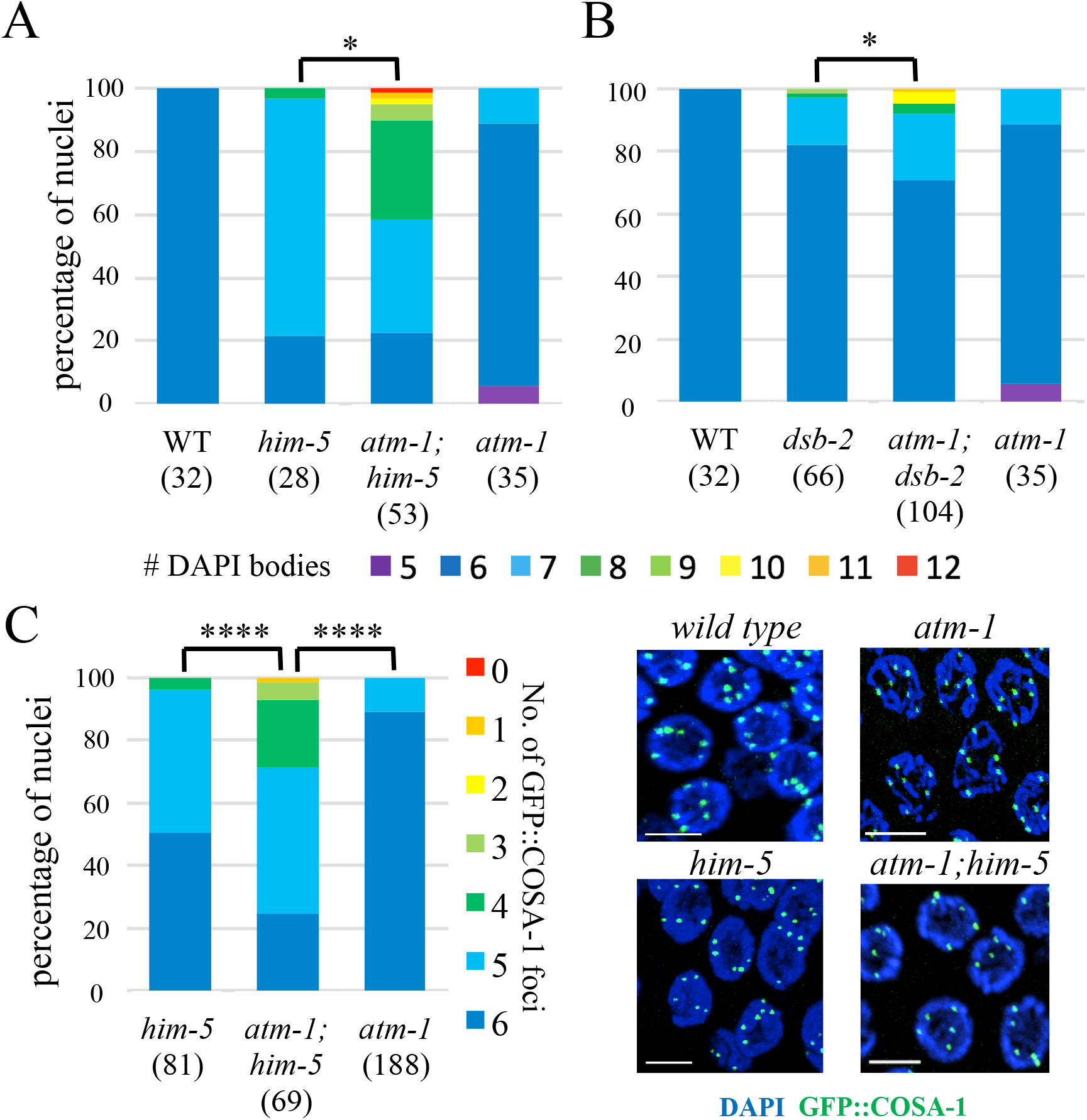
*atm-1* mutations exacerbate CO defects of DSB-defective mutants (A) *him-5* (B) *dsb-2.* Quantification of DAPI bodies in −1 nuclei. *P<0.05, two-tailed Mann Whitney test. (C) Proportion of nuclei with indicated numbers of GFP::COSA-1 foci ****P<0.0001, two-tailed Mann Whitney test. Three gonads were analyzed for each genotype. Numbers below each genotype indicate the numbers of nuclei analyzed. (D) Images of *him-5* and *atm-1;him-5* late pachytene nuclei are shown with DAPI (blue) and GFP::COSA-1 (green). Scale bar = 5 μm.

The appearance of these DAPI-bodies was similar to *him-5*—with five well-formed bivalents and 2 uniformly-sized univalents—which would explain the 2-3% HIM phenotype in the homozygous stocks. When *atm-1* mutations were combined with *him-5* or *dsb-2*, the percentage of univalents chromosomes was significantly increased. This was most striking in *atm-1;him-5* where >40% of diakinesis-stage nuclei contain >8 DAPI bodies, the equivalent of two or more chromosomes with defective COs. Thus, while *atm-1* mutant animals have been reported to exhibit an increase in RAD-51 foci (Checchi *et al.* 2014), fewer COs appeared to form.

To validate these results, we also quantified COs using GFP::COSA-1, a fusion protein that localizes to the chiasma formed between homologs (Yokoo *et al.* 2012). Since worm chromosomes usually only receive a single CO, GFP::COSA-1 is seen as a single focus per homolog pair starting in mid-late pachytene. As reported previously, most *him-5* mutant nuclei have only 5 GFP::COSA-1 foci (Machovina *et al.* 2016) reflecting the loss of X chromosome CO formation (Figure 1C). Similarly, ~10% of nuclei in *atm-1* mutants contained only 5 GFP::COSA-1 foci in accord with fraction of diakinesis-stage nuclei that contain 7 DAPI bodies in this mutant (Figure 1C). The number of GFP::COSA-1 foci was also significantly reduced in *atm-1;him-5* compared to *him-5* (Figure 1C, P<0.0001). Together these data implicate *atm-1* as a pro-CO factor in *C. elegans.*

We further tested the impact of *atm-1* loss on CO formation by examining recombination rates. We examined recombination rates in two large regions of chromosome III, the intervals between *dpy-18,unc-64* which comprises >13cM of the right arm of the chromosome and between *dpy-1,lon-1* which spans ~14cM from the middle of the left arm into the middle of the central gene cluster. In both intervals, no significant change in genetic distance was found in *atm-1* single mutants compared to wild type (Table 1A, 1B). However, *atm-1;him-5* mutants gave a lower recombination rate compared to *him-5* mutants in both regions of chromosome III (Table 1A, 1B). These data support the interpretation that *atm-1* mutations exacerbate the recombination defects caused by lack of *him-5* function. Together, these data strongly argue that COs are reduced in *atm-1;him-5* double mutants.

**TABLE 1.** *ATM-1* MUTANTS EXACERBATE DEFECTS IN RECOMBINATION

| Genotype | N2 | *atm-1* | *him-5* | *atm-1;him-5* |
| --- | --- | --- | --- | --- |
| *dpy-18* (III: 8.85), *unc-64* (III: 21.20) |  |  |  |  |
| Recombination rate | 12.01% (1896) | 11.97% (2816) | 14.22%<sup>#</sup>(2475) | 10.93%<sup>***</sup>(1587) |
| *dpy-1* (III: -15.66), *lon-1* (III: -1.63) |  |  |  |  |
| Recombination rate | 17.56% (3995) | 15.99% (1958) | 17.14%<sup>***</sup> (2705) | 13.78%<sup>***</sup> (1652) |

Chi square test, <sup>#</sup>N2 vs. *him-5*, $p < 0.05$ ; <sup>\*\*\*</sup>*him-5* vs. *atm-1;him-5*, $p < 0.0001$ . All other comparisons, n.s.

### ATM-1 limits the number of early repair intermediates

To understand the nature of the increased RAD-51 foci in *atm-1* mutants (Checchi *et al.* 2014) but the decreased numbers of COs in *atm-1;him-5* and *atm-1;dsb-2* mutants, we considered the possibility that ATM-1 might have different roles in otherwise wild-type versus CO-limiting situations. In this scenario, ATM-1 might limit DSBs under normal conditions (leading to the observed increase in RAD-51 foci), but when confronted with sub-threshold COs (COs on some chromosomes, but not all), might function in a regulatory feedback loop to retain DSB activity (leading to reduced RAD-51 signals and fewer COs in the mutants). To determine if these different scenarios exist, we quantified RAD-51 foci in *atm-1* mutants. We note that prior studies showed a >95% concordance between RAD-51 foci and free ends marked by TUNEL staining, so that the former can be used as a surrogate to assess DSB levels (Mets and Meyer 2009). We first validated the results from the prior studies (Checchi *et al.* 2014): using a different source of RAD-51 antibodies (See Materials and Methods), we also observed an increase in RAD-51 foci in *atm-1* mutant germ lines compared to wild type (P<0.0001; Chi-square test; Figure 2B-D, S1A and S1B). In *atm-1;him-5* and *atm-1;dsb-2* double mutants, distribution of RAD-51 foci in the pachytene germ line was altered compared to either single mutants (p<0.0001; Chi-square test; Figure 2D-I). Average RAD-51 levels were also elevated in *atm-1;him-5* compared to *him-5* (p=0.03; Wilcoxon matched-pairs signed rank test, Figure 2F). These differences are particularly striking in light of our observations that COs were decreased in both *atm-1;him-5* and *atm-1;dsb-2* (Figure 1). Thus, we conclude that RAD-51 dynamics are affected by loss of *atm-1* function in wild type and in mutants where crossovers are limiting.

**Figure 2.**
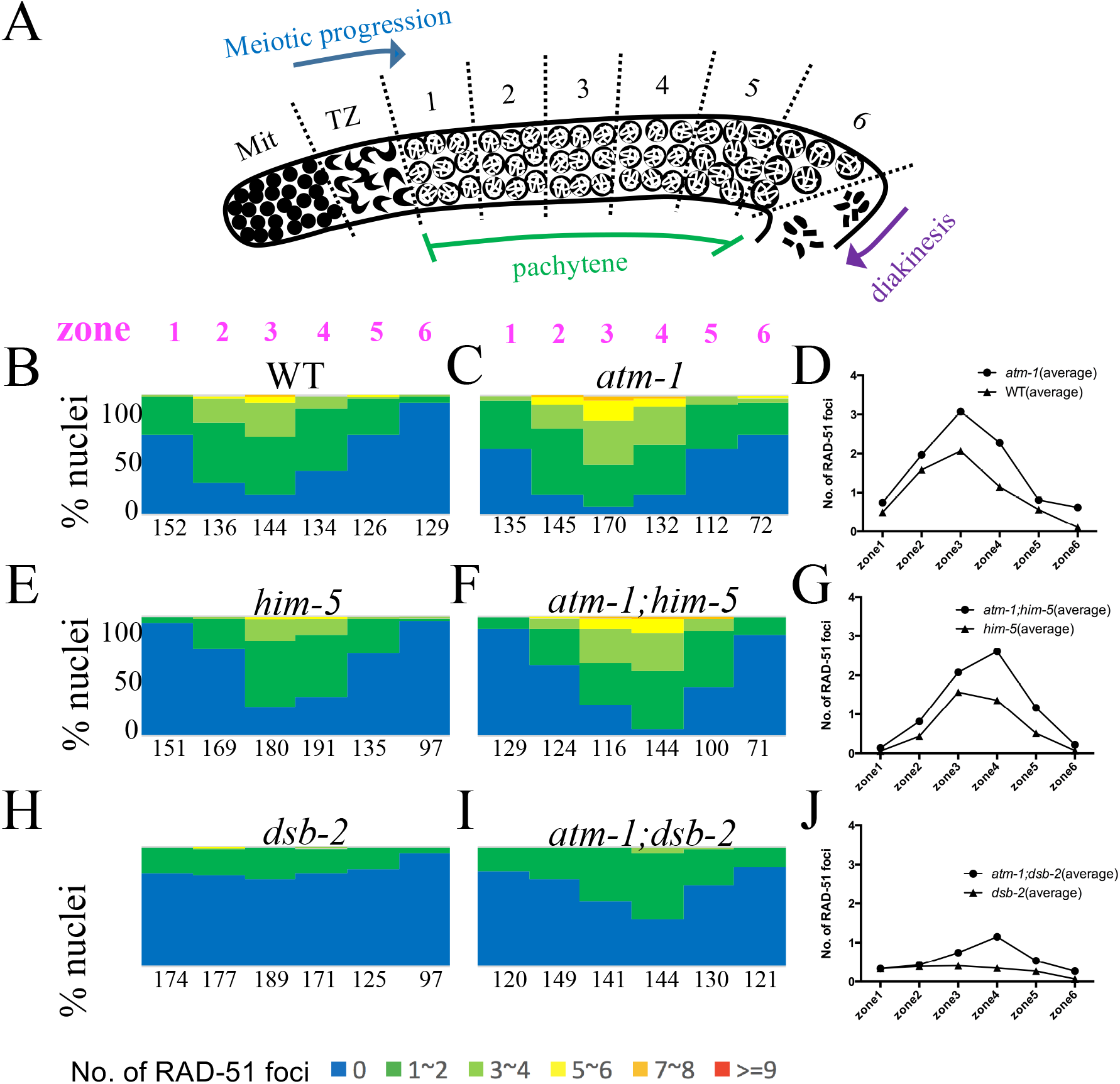
Early repair intermediates accumulate to a greater extent in *atm-1* mutants. (A) Schematic of the *C. elegans* germline showing regions in which RAD-51 foci were quantified. Mit: Mitotic zone. TZ: transition zone. (B, C, E, F, H, I) The percentage of nuclei in each zone containing the indicated number of RAD-51 foci shown in the color key at the bottom. Numbers represent total # of nuclei counted/zone for three gonads/genotype. (D, G, J) Comparison of the average number of RAD-51 foci for each genotype.. # = nuclei scored/ 3 germ lines/ genotype. χ^square^, *atm-1* vs. N2; *atm-1;him-5* vs. *him-5*; and *atm-1;dsb-2* vs. *dsb-2*: p<0.0001 for each. Representative images of germ lines are shown in Supplemental Figure S1.

RAD-51 accumulates on single-stranded, resected DNA ends as a filament that is dismantled upon stable strand exchange. Excessive RAD-51 signal would be seen if additional DSBs were present or if the kinetics of RAD-51 filament formation or turnover were altered. To distinguish between these possibilities, we quantified RAD-51 foci in *rad-54* mutant animals in which strand invasion cannot occur and therefore RAD-51 filaments accumulate. As expected, we observed increased RAD-51 foci in *atm-1 rad-54* single mutants compared to *rad-54* (Table 2 and (Checchi *et al.* 2014)). We also see more RAD-51 foci in *atm-1,rad-54;him-5* mutants compared to *rad-54;him-5* (Table 2). In both wild-type and *him-5* mutants, loss of *atm-1* resulted in ~5 additional DSBs.

**TABLE 2.** EARLY REPAIR INTERMEDIATES ACCUMULATE IN *ATM-1* MUTANTS

| genotype | No. of RAD-51 foci/nucleus |
| --- | --- |
| *rad-54* | 31.08±1.88 |
| *atm-1;rad-54* | 35.91±1.64<sup>§</sup> |
| *rad-54;him-5* | 22.83±2.35 |
| *atm-1;rad-54;him-5* | 28.82±3.73<sup>†</sup> |

Analysis of late pachytene nuclei
<sup>§</sup>t test, $p < 0.0001$
<sup>†</sup>t test, $p < 0.0001$

In addition to SPO-11 mediated DSBs, unrepaired mitotic or meiotic S phase DNA damage can contribute to pachytene accumulation of RAD-51. To determine if such damage is present in *atm-1* mutants, we analyzed RAD-51 foci in *spo-11* and *atm-1*; *spo-11* in which meiotic DSBs are not formed. If pre-meiotic (or meiotic S-phase) damage were carried through into pachytene, RAD-51 foci should be more prevalent in *atm-1;spo-11* compared to *spo-11.* We previously showed that ~10% of *spo-11* nuclei have GFP::COSA-1 foci and elicit CO feedback mechanisms (Machovina *et al.* 2016). This result was supported here by the appearance of a small number of RAD-51 foci in the pachytene region of *spo-11* mutants. By contrast, we saw very few RAD-51 foci in the pre-meiotic and meiotic regions *of atm-1;spo-11* (Figure 3A, 3F, S1C and S1D). These results suggest that the extra RAD-51 foci in *atm-1* pachytene nuclei are not a consequence of pre-meiotic damage. Instead, these results point to a role for ATM-1 in limiting meiotic DSB formation and/or the early processing of these DSBs.

**Figure 3.**
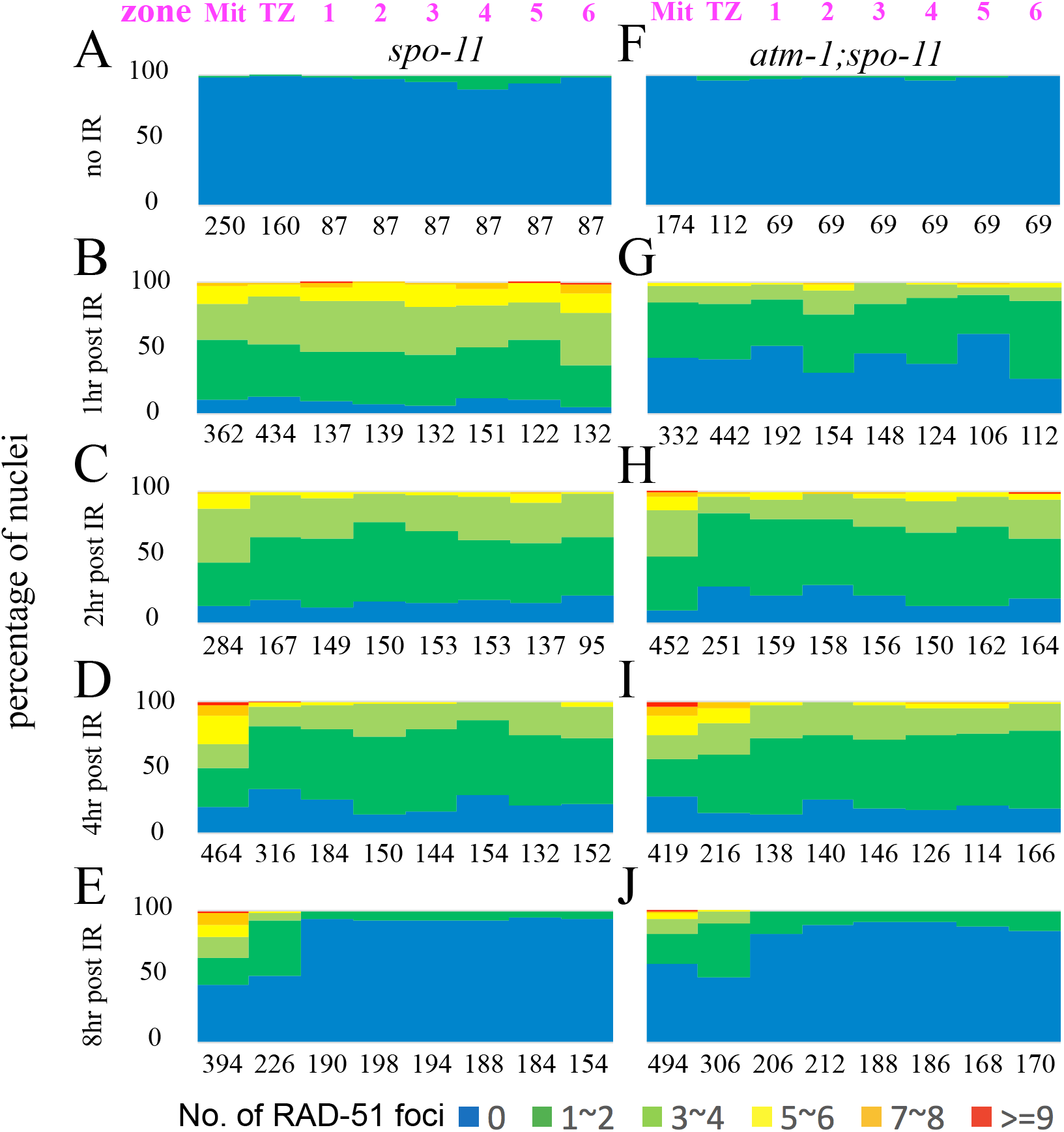
Loading of RAD-51 is delayed in *atm-1* mutants. For each region, the percentage of nuclei with a given number of RAD-51 foci are shown as a heat map from 0 - >9 as indicated below. Zones are shown in Figure 2A. # = nuclei scored/ 3 germ lines/ genotype. (A-E) *spo-11* mutants. (F-J) *atm-1;spo-11* mutants. Shown in a time-course post-exposure to lOGy IR (A, F): unirradiated controls (B, G): 1 hour post-exposure shows reduced loading in *atm-1;spo-11.* (C, H): 2 hours and (D, I): 4 hours post-exposure loading is almost indistinguishable between control and *atm-1;spo-11.* (E, J): 8 hours post-exposure, most RAD-51 foci have been removed in the meiotic region of the germ line. We note that repair in the mitotic region is resolved with distinct kinetics.

### RAD-51 loading is impaired in *atm-1* mutant animals

We next set out to determine if *atm-1* mutants might be defective in RAD-51 filament formation and/or processing. Since the appearance and disappearance of RAD-51 foci is influenced both by the extent of SPO-11 activity and the kinetics of RAD-51 loading and disassembly, it can be difficult to tease out the impact of altered DSB dynamics from altered behavior of RAD-51. To specifically examine the requirement for ATM-1 in RAD-51 loading, we therefore assayed RAD-51 focus formation in backgrounds where meiotic breaks are not made, comparing *atm-1;spo-11* with *spo-11* at various time points after exposure to IR. Surprisingly, at 1 hour-post IR, we observed many fewer RAD-51 foci in *atm-1;spo-11* double mutants compared to *spo-11* (Figure 3B, 3G, S1E and S1F). These results intimate that early RAD-51 accumulation is impaired in *atm-1* mutants. At 2 hours-post IR, the number of RAD-51 foci in *atm-1;spo-11* reached levels comparable to *spo-11* 1-hour-post IR (Figure 2C, 2H, S1G and S1H). RAD-51 foci in *atm-1;spo-11* and *spo-11* decreased comparably as shown by the RAD-51 signals at 4 and 8 hrs post-IR (Figure 2D, 2E, 2I and 2J). Thus, while RAD-51 focus formation may be affected by loss of *atm-1*, processing/removal of RAD-51 appears to be normal. While IR-induced and SPO-11 induced breaks are not treated identically in cells (Macaisne *et al.* 2018), these data raise the possibility that the excess RAD-51 foci seen in *atm-1, atm-1;him-5* and *atm-1;dsb-2* are not due to impaired RAD-51 processing, but rather most likely result from additional meiotic DSBs.

### ATM-1 inhibits homolog-independent repair, channeling DSBs towards inter-homolog recombination

The bulk of meiotic DSBs are repaired through homologous recombination using either IH-HR or IS-HR repair pathways, and among these two pathways, only IH-HR can create chiasmata. We hypothesized that ATM-1 promotes CO formation by inhibiting processes that do not engage the homolog, thus channeling more DSBs going through IH-HR. Without this inhibition, DSBs would preferentially be repaired through IS-HR or as noncrossovers, leading to a deficit in COs. To test this hypothesis, we took advantage of two mutations that are known to impair homolog-independent repair: *brc-1* and *smc-5* (Adamo *et al.* 2008; Bickel *et al.* 2010). Mutants carrying either mutation had a small percentage of diakinesis-stage nuclei with chromosome fragments. In *syp* mutants where synaptonemal complex (SC) formation is impaired, the homolog is not readily available for HR and IS-HR is thought to be the major DSB repair pathway (Adamo *et al.* 2008; Bickel *et al.* 2010; Macaisne *et al.* 2018). In the *syp* mutant background, both *brc-1* and *smc-5* showed increased chromosome fragmentation. In contrast to *brc-1;syp-1* and *smc-5;syp-1*, we did not observe chromosome fragments in *atm-1;syp-1* mutants (Figure 4) indicating that all DSBs are repaired in this mutant context. By contrast, in *atm-1;brc-1* mutants, we found more nuclei with fragments compared to *brc-1* mutants (Figure 4, P<0.05). A similar result was observed in *atm-1;smc-5* mutants, in which ~50% more nuclei contained DNA fragments compared to the *smc-5* single mutant. Further, *atm-1;smc-5;brc-1* triple mutants revealed a substantially higher percentages of nuclei with fragments compared to *smc-5;brc-1* double mutants (Figure 4).

**Figure 4.**
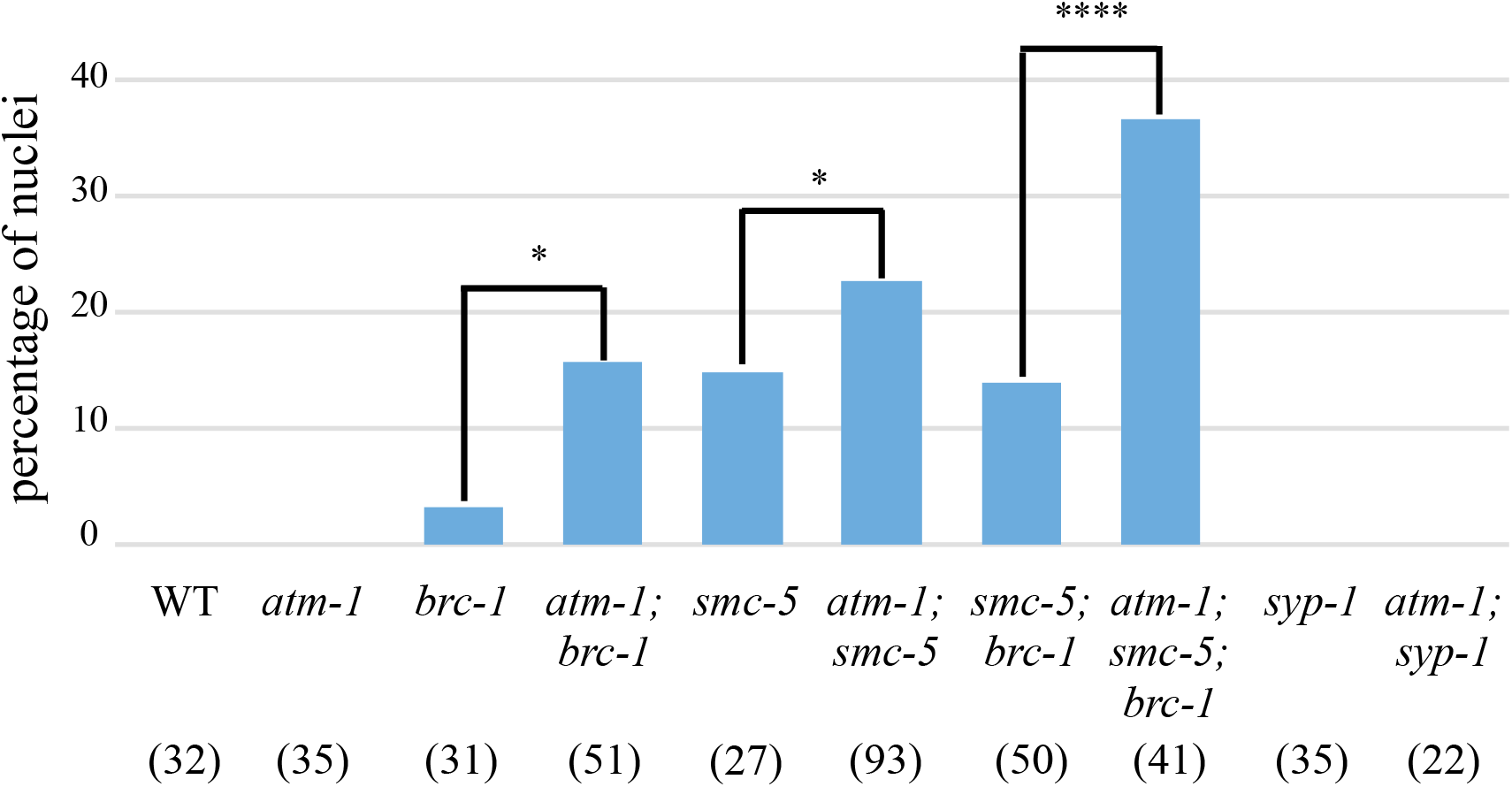
*atm-1* mutation increases fragmentation when inter-sister repair is impaired. Percentage of-1 nuclei containing DNA fragments as seen by DAPI staining. Numbers in brackets are the number of nuclei analyzed/ genotype. *P<0.05, ****P<0.0001 (χ^2^ test).

To determine if the increased number of chromosome fragments in the *atm-1* mutants arises simply as a result of excess meiotic DSBs formed in this background, we reasoned that *atm-1* should have no impact on the repair of DSBs induced by IR in the *smc-5;spo-11(-)* background. Consistent with our results in *spo-11*(+), more nuclei with fragments were observed in *atm-1;smc-5;spo-11* mutants compared to *smc-5;spo-11* after exposure to 10 Gy IR, while a similar percentage of nuclei with fragments was apparent in non-irradiated *smc-5;spo-11* and *atm-1;smc-5;spo-11* (Figure S2). Together, these data suggest that ATM-1 functions outside of DSB break formation to promoting homolog-dependent repair.

### ATM-1 functions on SPO-11 dependent and independent DSBs to promote CO formation when the number of DSBs is limiting

The CO defects in *atm-1;him-5* and *atm-1;dsb-2* appeared stronger than expected based on the mild CO defect in *atm-1* single mutants. This led us to investigate whether *atm-1* differentially affects DSB outcomes under low DSB versus high DSB situations. We compared CO outcomes in *spo-11* and *atm-1;spo-11* mutants and after exposure to 2 Gy, 10 Gy, and 25 Gy IR (Machovina *et al.* 2016). Upon exposure to 2 Gy or 10 Gy IR, nuclei of *atm-1;spo-11* double mutants exhibited greater numbers of DAPI bodies (fewer bivalents) at diakinesis compared to *spo-11* single mutants (Figure 5A, S3). The impact on COs is not specific to the *spo-11* mutant background as fewer bivalents were also seen in irradiated *atm-1;dsb-1* compared to *dsb-1* mutant animals that are also defective in meiotic DSB formation (Stamper *et al.* 2013) (Figure 5A, S3). By contrast, no difference in CO outcomes was observed when *atm-1* and *atm-1;spo-11* mutants were exposed to 25 Gy, both mostly contained 6 DAPI bodies at diakinesis (Figure 5B). Since 1 Gy IR is expected to give ~2 meiotic DSBs (Machovina *et al.* 2016) and to increase linearly with dose, these results suggests that a threshold exists somewhere between 20 and 50 DSBs beyond which *atm-1* dysfunction in DSB repair is overcome.

**Figure 5.**
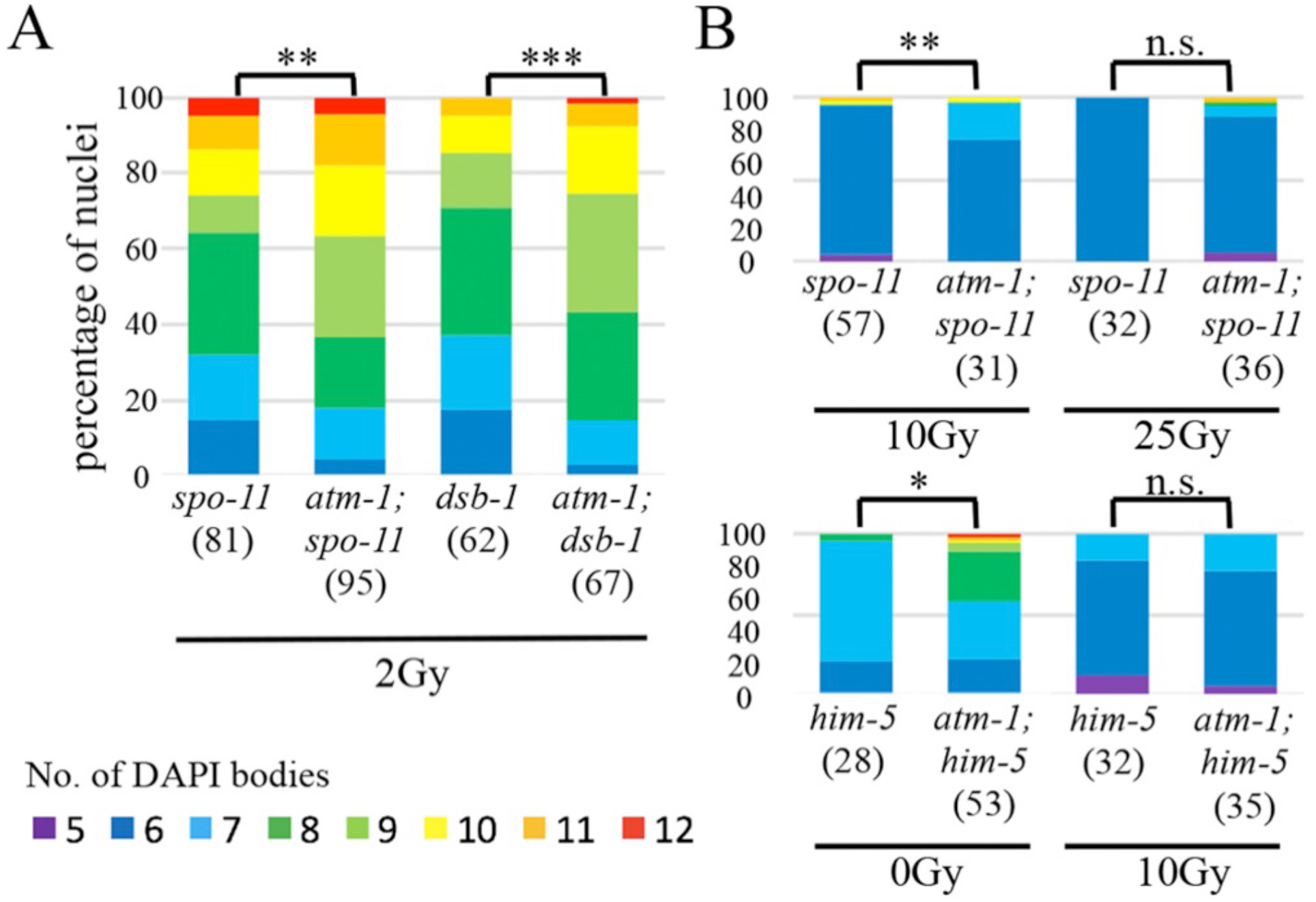
*atm-1* acts on exogenous DSBs as well as SPO-11 induced DSBS to promote crossovers within a threshold of total DSBs. Quantification of DAPI bodies in −1 nuclei for different doses of IR. Color key below. (A) 2 Gy IR leads to more COs in *atm-1;spo-11* and *atm-1;dsb-1* compared to control *spo-11 dsb-1*, respectively. (B, C) *atm-1* loss is overpowered by high numbers of DSBs, between 10 Gy and 25 Gy (20-50 DSBs) in *atm-1; spo-11* (top); below 10 Gy (20 additional breaks) in *atm-1;him-5* (~28 endogenous breaks). *P<0.05, **P<0.01, ***P<0.001, n.s.= no significant difference, two-tailed Mann Whitney test.

Our analysis of *atm-1;him-5* mutants provided support for a threshold of DSBs for ATM-1 regulation. As discussed above, in *atm-1;him-5* double mutants, DSBs appeared to be shunted into noncrossover repair pathways. However, the addition of 10Gy in this genetic background was sufficient to drive DSBs into the IH-HR pathways, leading to 6 bivalents in most diakinesis nuclei (Figure 5B). The exposure of *atm-1;him-5* to 10 Gy IR would induce ~20 more DSBs, bringing break levels to near those in *atm-1;spo-11* + 25Gy. Together these results support a role for ATM-1 in repair pathway choice when the number of DSBs is under a threshold of ~30-50 DSBs. When above this threshold, ATM-1 function appears to be bypassed.

### ATL-1 contributes to CO formation by influencing accumulation of early break intermediates

One candidate for subsuming ATM functions is ATR, the other major kinase involved in the DNA damage response. ATR is encoded by *atl-1* (ATM-Like) and is an essential gene that is required for mitotic DNA repair ((Garcia-Muse and Boulton 2005), reviewed in (Budzowska and Kanaar 2009)). Loss of *atl-1* leads to both macro- and micro-nuclei due to the impairment in DNA damage signaling (Figure S4A). In nuclei that go on to make oocytes, we observed nearly 25% with only 5 DAPI bodies (Figure 6A), presumably reflecting the formation of chromosome fusions in response to DNA damage. Another 5% of nuclei had more than 6 DAPI bodies (Figure 6A), suggesting that, like *atm-1, atl-1* is required for a full complement of CO exchanges. In *dsb-2* and *him-5* mutants, lack of ATL-1 function also reduced CO numbers (Figure 6A). Thus, we conclude that ATL-1, like ATM-1, contributes to CO formation when DSBs are limiting.

**Figure 6.**
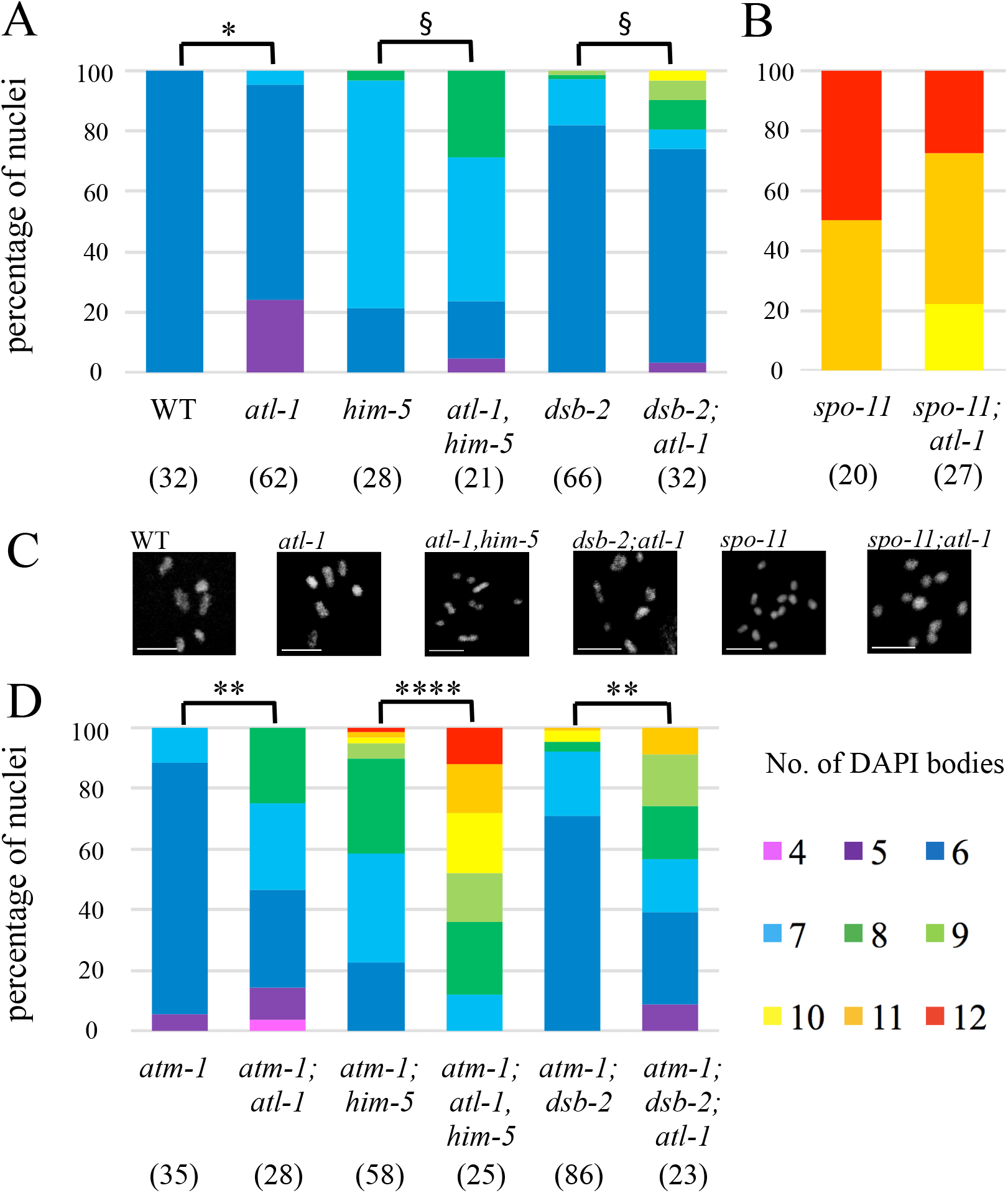
*atl-1* mutant animals exhibit defects in CO formation in wild-type and DSB-limiting situations. (A, B, D). Quantification of DAPI bodies in −1 nuclei containing the indicated number of DAPI bodies. *P<0.05, **P<0.01, ****P<0.0001, two-tailed Mann-Whitney test. § P<0.05, Fisher’s exact test. (A) *atl-1* has an increase in univalents alone or in combination with *him-5* and *dsb-2.* (B) *atl-1* mutants increase bivalents in *spo-11*, indicating a substantial amount of carry-though DNA damage from the mitosis and meiotic S phase. (C) Representative diakinesis-stage nuclei of different genotypes showing normal karyotype (wt) and mutant background with different proportions of bivalents and univalents. (D) Quantification of DAPI bodies in *atm-1;atl-1* double mutants alone or with DSB-defective mutations shows synergistic effects from the loss of both gene functions.

To determine if ATL-1 directly impacts DSBs/early repair processing, we analyzed RAD-51 dynamics. Two aspects of RAD-51 accumulation distinguished *atl-1* mutants from wild type. First, the appearance of RAD-51 signals differed: a subset of *atl-1* mutant nuclei exhibited very high RAD-51 signals (Figure 7A-C). Since the RAD-51 signal was so extensive in some nuclei, we quantified these images by binning the nuclei based on number of foci that were observed (Figure 7H-J). The second distinct aspect of *atl-1* mutants was the timing of RAD-51 focus formation: in *atl-1*, RAD-51 foci were observed in most nuclei from the transition zone (TZ) through the pachytene-diplotene border in *atl-1* (Figures 7C, H); whereas in wild type, foci only started to accumulate in the TZ and were seen in most nuclei only at mid-pachytene (Figures 7B, 7H, 2B, S1A).

**Figure 7.**
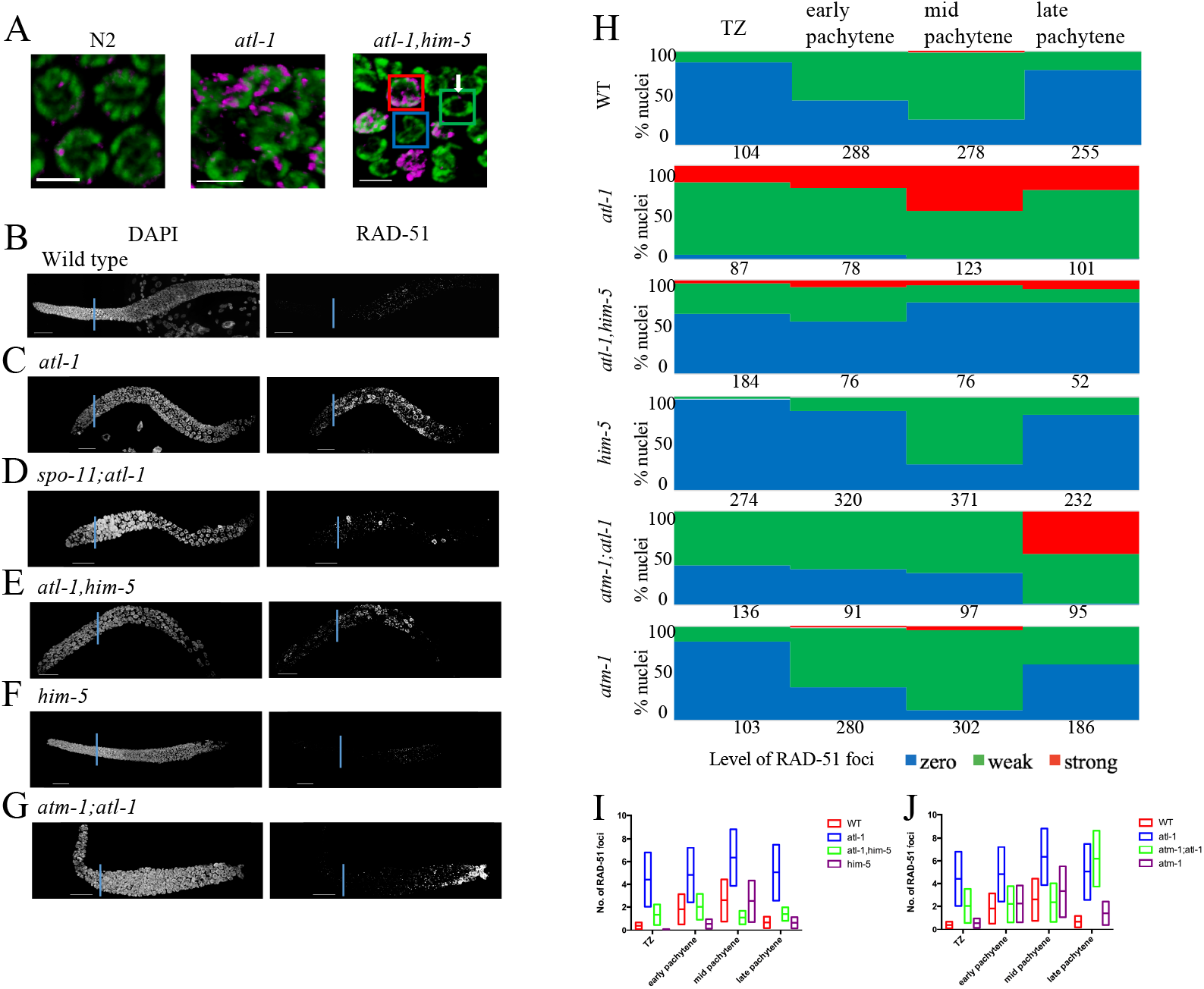
RAD-51 loading is altered by *atl-1*. (A) RAD-51 staining in pachytene nuclei of wild type N2, *atl-1*, and *atl-1 him-5.* RAD-51 (magenta); DAPI (green). Scale bar = 5 μm. The squares represent the three class of nuclei quantified in Figure 7C: Zero RAD-51 foci (blue); Weak staining (green); strong staining (red). (B-G) Representative images of DAPI and RAD-51 stained gonads from one-day old adults of indicated genotype. (H) Proportions of nuclei containing zero (blue), 1-6 (green, weak), or >6 foci (red, strong) RAD-51 foci in the leptotene-pachytene regions of the germ line as described in Materials and Methods. Numbers indicate total number of nuclei counted for each region for at least 3 germ lines/ genotype. (I, J) Range and average of RAD-51 foci for germ lines quantified in (H) for shown genotypes.

The differences in RAD-51 accumulation, together with the known role for *atl-1* in mitotic cell divisions (Abraham 2001; Garcia-Muse and Boulton 2005; Lawrence *et al.* 2015), led us to explore whether any of RAD-51 foci could be explained by an increase in carry-through damage from mitotic divisions. In *spo-11;atl-1*, there was substantial RAD-51 signal in the pachytene region indicating the presence DNA damage that arose independently from meiotic-induced DSBs (Figure 7D; compare with *spo-11* in Figure S1C). We note that this damage can contribute to CO formation as we saw an increase in bivalents in *spo-11;atl-1* diakinesis-stage oocytes (Figure 6B). Irradiation of *spo-11;atl-1* mutants with 10 Gy IR showed efficient RAD-51 loading within 1 hour post-IR and, as in the *atl-1* background, a subset of the signal appeared in stretches (as opposed to smaller foci; Figure S4D). Since these stretches were not seen in the irradiated *spo-11* mutant germ lines (Figure S1E, G), we infer that these stretches do not result from two adjacent DSBs, but rather from a defect that is specific to the *atl-1* mutant. Similar stretches of RAD-51 have been seen in meiotic mutants with defects in RAD-51 filament maturation (Mets and Meyer 2009; Ward *et al.* 2010), raising the possibility that ATL-1 limits/antagonizes RAD-51 loading. *atl-1* loss would therefore lead to excessive and perhaps, untimely, RAD-51 filament formation.

We also observed RAD-51 foci in the TZ of *atl-1,him-5* double mutant animals (Figure 7A, E). In this mutant background, the kinetics of RAD-51 formation and loss are similar to *atl-1* single mutants, appearing earlier and brighter that in wild type (Figures 7H). However, the number of nuclei with RAD-51 staining was reduced in *atl-1,him-5* compared to *atl-1* mutants, as expected since *him-5* mutations partially impair DSB formation. The number of RAD-51-positive nuclei was also reduced compared to *him-5* (Figure 7E-F, 7H-I): whereas upwards of 70% of *him-5* mutant nuclei stained weakly for RAD-51 at its peak, fewer than 30% of *atl-1,him-5* nuclei showed RAD-51 foci (Figure 7H-I). Of those that had foci, a subset had the very strong RAD-51 signals that were associated with carry-through damage, described above. Thus, 30% is an overrepresentation of the number of nuclei with *bona fide* meiotic damage. These data show that *atl-1* and *him-5* both contribute to RAD-51-focus formation and that *atl-1* is epistatic to *him-5* for timing of RAD-51 formation.

In yeast and mice, ATM and ATR function together to establish DSB homeostasis (Carballo *et al.* 2013): ATM inhibiting DSBs to prevent over-accumulation; ATR promoting DSBs to ensure sufficient COs can be made. We therefore wanted to know what the impact on meiotic CO formation would be when both *atm-1* and *atl-1* are mutated. To our surprise, *atm-1;atl-1* mutants showed a severe defect in CO formation: whereas both *atm-1* and *atl-1* single mutants exhibited mild CO defects (Figure 6A and 6C), in the double mutants, >70% of nuclei had CO defects, of which >50% contain one or more pairs of univalents. Loss of both *atm-1* and *atl-1* also exacerbated the CO defect associated with *him-5* and *dsb-2* (Figure 6C). Thus, despite seemingly antagonistic roles on RAD-51 formation, *atm-1;atl-1* double mutants are significantly impaired in CO formation.

Consistent with these results, we saw very few RAD-51 foci in early and mid-pachytene nuclei in *atm-1;atl-1* (Figures 7G-H, J). At late pachytene, almost all nuclei stained strongly for RAD-51, a phenotype seen in neither of the single mutants. It is unlikely that all of these nuclei are destined for apoptosis since we observed diakinesis-stage oocytes with well-formed bivalents (Figure 6C). Instead this data suggests a change in CO regulation in the double mutant (discussed below).

### CO feedback is impacted by loss of *atm-1* and *atl-1*

Defects in CO formation are thought to activate surveillance systems that feed back onto the break machinery to maintain DSB competency when COs are not detected (Rosu *et al.* 2013; Stamper *et al.* 2013; Machovina *et al.* 2016; Nadarajan *et al.* 2017; Pattabiraman *et al.* 2017). The activity of this feedback mechanism is spatially observed in the pachytene germ line as an extended region of DSB-1 and DSB-2 staining in CO-deficient worms (Rosu *et al.* 2013; Stamper *et al.* 2013). We reasoned that the increased number of RAD-51 foci observed in *atm-1* mutants might be explained by changes in DSB-2 regulation. We observed that DSB-2 staining in *atm-1* mutants was shifted proximally—turning on slightly later than in wild type relative to the onset of leptotene (Figure 8). It also persisted slightly longer, taking on average ~7% more of the pachytene region than wild type (Figure 8). The small number of excess DSBs in *atm-1* may be attributed to this increased window of opportunity for DSB-2- (and by inference, DSB-1-) dependent breaks. We also observed that *atm-1* activity was not required to induce (or maintain) the extended domain of DSB-2 in *him-5* mutants (Figure 8). Thus, we posit that the reduction in COs in *atm-1;him-5* cannot be explained by inhibition of DSB-2.

**Figure 8.**
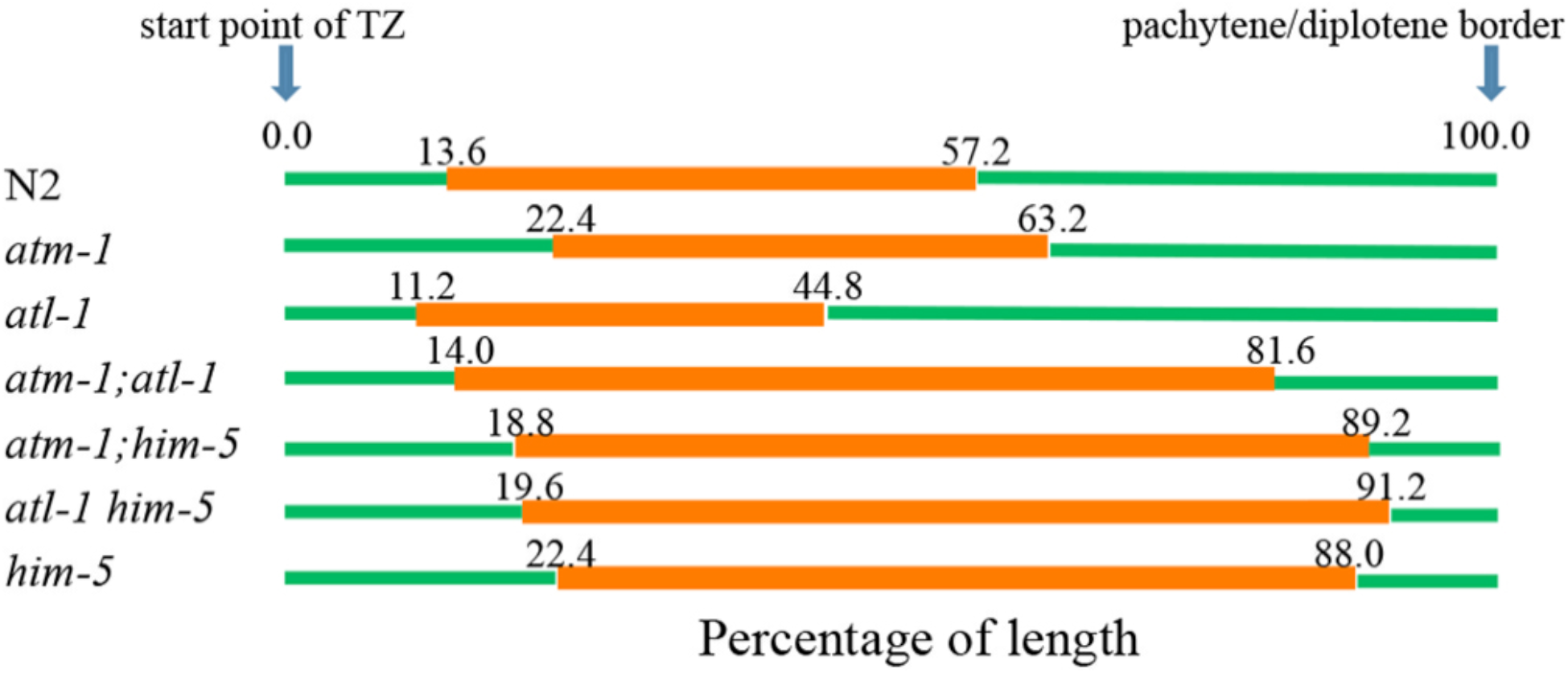
DSB-2 staining manifests different kinetics in the absence of ATM-1 or ATL-1 functions. Proportion of length of the germline region from meiotic onset to the pachytene/diplotene border that stain positive for nuclear localized DSB-2. Non-stained regions (green); stained (red). Genotypes are shown to the left with the average DSB-2 region depicted in orange (n= 3, 6, 7, 4, 4, 5, 6 for N2, *atm-1, atl-1, atm-1;atl-1, atm-1;him-5, atl-1 him-5*, and *him-5*, respectively).

In *atl-1* mutants, DSB-2 staining spanned only ~33% of the leptotene to pachytene region whereas it comprised ~45% in wild type (Figure 8). This result suggests that the CO-dependent deactivation of DSB-2 occurred more rapidly in *atl-1* mutants. *him-5* is epistatic to *atl-1*, as was seen by the slight delay in DSB-2 onset and much extended domain of staining in *atl-1,him-5* double mutants. These phenotypes are best explained by HIM-5’s role in promoting DSB formation: delaying the formation of DSBs and preventing DSBs on the X chromosome (Meneely *et al.* 2012).

In the *atm-1;atl-1* mutants, the DSB-2 region was distinct from either single mutant: DSB-2 was activated as in wild type, yet it persisted until very late pachytene. The extension of DSB-2 staining was similar to that seen in *him-5* mutants, suggesting it may also reflect the diminished COs that form in the double mutants.

## DISCUSSION

ATM-1 and ATR-1 have both unique and overlapping functions in worm meiosis that influence the formation of COs. We have shown that these genes have antagonistic and synergistic roles in DSB and CO formation which is summarize in our proposed model in Figure 9. We posit that upon SPO-11 activation, a small number of initial DSBs are made (<10) that are sufficient to activate ATM-1 and ATL-1. Both proteins would then influence the activation of DSB-2 (perhaps through DSB-1): ATL-1 directly and ATM-1 through the regulation of early events post-DSB formation, perhaps resection. The lack of input from ATL-1 would explain the reduction in DSBs seen in *atl-1* mutants. Delayed resection could explain the delayed activation of the DSB-2 feedback loop and its prolonged localization in *atm-1* mutants. We propose that ATM-1 and ATL-1 then influence the transition from resection to an IH-CO competent RAD-51 filament. In the case of ATM-1, our *atm-1;spo-11 + IR* data predict a role in the timely recruitment of RAD-51. This delay could be a consequence of its role in resection or its role in influencing RAD-51 loading. Further studies will be required to determine if ATM-1 has the same impact on RAD-51 recruitment at SPO-11-induced breaks.

**Figure 9.**
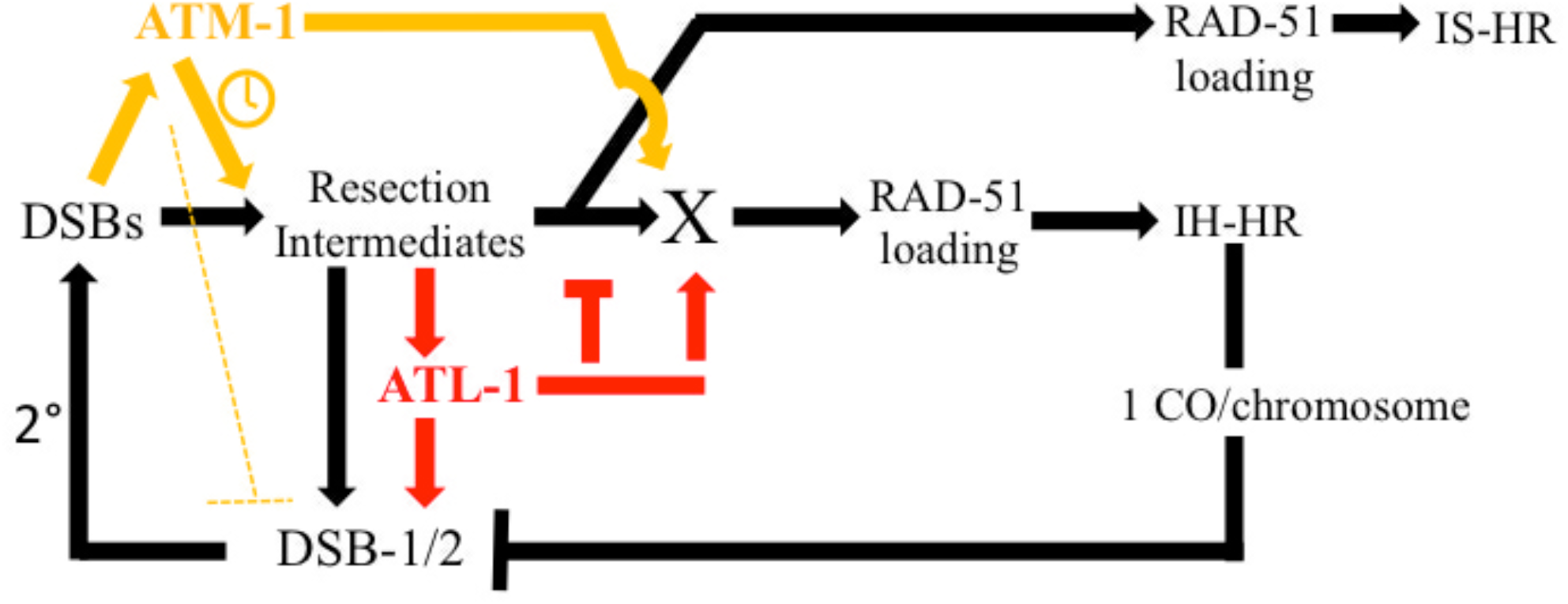
Model for ATM-1 and ATL-1 functions during meiosis. We propose that ATM-1 is activated by DSBs and then promotes the timely resection of ends and repair using the homolog as a repair template. The excess RAD-51 foci in *atm-1* mutants could be explained if ATM-1 inhibits DSBs (dotted line), perhaps through DSB-1, but possibly SPO-11 or other accessory factors. Alternatively, delayed resection and IH-CO formation could keep the feed-forward loop active longer to allow additional DSBs to be made. We further posit that ATL-1 is activated by resection intermediates and both ATL-1 and a subset of pre-RAD-51 repair intermediates feed forward to promote additional DSB formation. ATL-1 then inhibits extensive resection and redundantly tp ATM-1 helps to promote RAD-51 loading. These two activities can explain the delayed, excessive RAD-51 loading in *atm-1;atl-1* double mutants.

To our surprise, we found that despite increased RAD-51 foci, COs are diminished in *atm-1* mutants. This reduction could due to the ability of ATM-1 to influence the timely recruitment of RAD-51 (assuming SPO-11 induced breaks are treated the same at IR-induced breaks) and/or to shunt meiotic DSBs into the IH-CO repair pathway. In its absence, noncrossover repair pathways are favored. Thus, *atm-1* appears to function antagonistically on different aspects of CO formation, limiting the total number of DSBs, but increasing the likelihood that DSBs are repaired by IH-CO repair. ATL-1 also appears to have an antagonistic function with decreased numbers of DSBs yet excessive RAD-51 loading seen in the *atl-1* mutant animals. These mutually antagonistic behaviors can be explained by negative and positive feedback loops and built-in functional redundancy (Figure 9) that illustrate the extensive and precise machinations required to ensure that COs occur on each chromosome.

Our model posits that, as in other systems (reviewed in (Moriel-Carretero *et al.* 2018)), DSBs activate ATM-1 to promote timely resection and subsequent activation of ATL-1. Resected ends and ATL-1 would both promote a secondary wave of DSBs that are induced through activation of DSB-2, and by inference DSB-1 (Stamper *et al.* 2013). This explains the decreased RAD-51 foci in *atl-1* mutants and the delay in DSB-2 loading in *atm-1* mutants. The increased number of RAD-51 foci in *atm-1* mutants might suggest that ATM-1 negatively regulates DSB formation. Alternatively, the 5-6 extra DSBs (Table 2) that are made could also be explained by persistent activation of the CO surveillance system ((Machovina *et al.* 2016), Figure 9). In this case, a bias for IS-HR or noncrossovers in *atm-1* mutants would result in later CO formation and tardy deactivation of DSB-1/2. ATL-1 (and perhaps ATM-1) are likely to be regulating DSBs and DSB-2 localization through phosphorylation of SPO-11 and/or one or more its accessory factors. Bioinformatic analysis identified potential (S/T)Q sites in all of the ten known factors that influence DSB formation supporting the possibility that one or more are direct targets of ATL-1 (or ATM-1) (Fig S5). Our analyses of *atm-1;dsb-2* and *atm-1;him-5* double mutants showed increased DSBs (seen as RAD-51 foci), ruling out DSB-2 and HIM-5 as the sole targets of ATM/ATR signaling, although they may have redundant functions. Further studies are needed to identify the relevant targets in the DSB machinery.

Our observation that RAD-51 loading is delayed in *atm-1* mutants post-IR suggests that ATM-1 facilitates efficient formation of the RAD-51 filament. In mitosis, both ATM and ATR activate resection activities with ATR functioning as well to attenuate Exo1-mediated activities (reviewed in (Gobbini *et al.* 2013)). In meiosis, recent studies have shown that yeast ATM1/Tel1 promotes resection of early DSBs (Joshi *et al.* 2015) and that mouse ATM helps to initiate and promote resection (Mimitou *et al.* 2017). A defect in resection could explain the delay in RAD-51 recruitment in *atm-1;spo-11* post-IR. A likely downstream target to regulate resection and RAD-51 loading (serving as protein X in Figure 9) is RAD-50, whose homologs are targets of ATM signaling in other systems (Gatei *et al.* 2011). In worms, RAD-51 loading in the early-mid pachytene region requires RAD-50 activity, whereas late pachytene loading is RAD-50-independent (Hayashi *et al.* 2007). The transition between these two states is thought to correspond to the switch from IH-HR to IS-HR. By promoting RAD-50 activity, ATM-1 could therefore assure the repair of meiotic DSBs by IH-HR; in its absence, RAD-50 activity would be attenuated and IS-HR/ noncrossover repair would be favored.

ATR has also been implicated in resection control, both promoting and restraining extensive EXO1 activity (Tomimatsu *et al.* 2017). Our observation that a subset of ATR mutants present with large RAD-51 aggregates intimates that the latter function, at least, of ATR may be conserved in worm ATL-1. The rapid disappearance of nuclear localized DSB-2 in *atl-1* mutants could be explained by the formation of longer resected in ends that would more rapidly be converted into IH-COs, leading to cessation of DSB formation by CO feedback (Machovina *et al.* 2016). Together with our analysis of ATM-1, these data raise the intriguing possibility that different resection tract lengths could influence the crossover vs noncrossover decision.

Based on their mutually antagonistic behavior on DSBs/RAD-51 foci formation, we explored the impact of loss of both ATM-1 and ATL-1 functions. In *atm-1;atl-1* double mutants, both RAD-51 foci and COs are decreased (Figure S4D and 6C). This suggests that *atl-1* has an additional role in promoting RAD-51 loading that is redundant with ATM-1. Although depicted as a shared target gene, this may reflect one or more downstream roles. We envision that in the double mutant background, the DSB feedback loop is activated via resected ends (through DSB-1 or other accessory factors), the ends are hyper-resected due to loss of *atl-1*, but RAD-51 loading is significantly delayed due to the joint functions of ATL-1 and ATM-1 on RAD-51 loading which are relieved when the block to IS-HR is relaxed in late pachytene, leading to the excessive loading of RAD-51 in the *atm-1;atl-1* double mutant.

These data reveal the complex interplay between ATM and ATR signaling in meiosis that ultimately helps determine both the number of DSBs and the formation of IH-COs. These studies highlight multiple potential targets of ATM-1 and ATL-1 and provide the basis for the future identification and analysis of specific substrates. These studies also illuminate the evolutionary conserved of antagonism between ATM and ATR that is necessitated by the requirement for CO formation on each chromosome.

---

Supplemental Table 1 List of strains generated for this study.

Supplemental Figure 1. ATM-1 limits the accumulation of RAD-51 foci.

Supplemental Figure 2. The impact of *atm-1* on noncrossover outcomes is independent of SPO-11 induced breaks.

Supplemental Figure 3. SPO-11-independent COs are induced in *atm-1* mutants, revealing carry-through damage from pre-meiotic events or meiotic S phase. At the same time, exposure to IR leads to fewer COs in *atm-1* mutant background, revealing a defect in converting DSBs to COs in the absence of ATM-1 function.

Supplemental Figure 4. RAD-51 analysis in *atl-1* mutants and post-IR exposure.

Supplemental Figure 5. Bioinformatic analysis of potential ATM-1/ATL-1 phosphorylation sites in *C. elegans* DSB proteins.

## ACKNOWLEDGMENTS

The authors wish to thank Dr. Kara Bernstein, Dr. Arjumand Ghazi, Dr. Arthur Levine, Dr. Nicolas Macaisne, Dr. Brooke McClendon, and Logan Russell for critical reading of the manuscript. Gratitude is extended to Dr. Sarit Smolikove for sharing her anti-RAD-51 antibody. Some strains were provided by the CGC, which is funded by NIH Office of Research Infrastructure Programs (P40 OD010440). W.L. was funded by Tsinghua University School of Medicine; J.L.Y. was funded by Pennsylvania Formula Funds and NIGMS/NIH (GM104007).

